# A *SUBTILISIN-LIKE SERINE PROTEASE1 (OsSUBSrP1)*, plays an important role in the wax and cutin pathway, is essential for panicle development in rice

**DOI:** 10.1101/2021.07.04.451081

**Authors:** Asif Ali, Tingkai Wu, Hongyu Zhang, Peizhou Xu, Syed Adeel Zafar, Yongxiang Liao, Xiaoqiong Chen, Yutong Liu, Wenming Wang, Xianjun Wu

## Abstract

Panicle degeneration is a severe physiological defect and causes reduction in grain yield. In this study, we characterized and presented the functional analysis of our previously reported mutant *apa1331* (*apical panicle abortion1331)* that showed apical spikelet degeneration. The anthers from the apical spikelets of *apa1331* were degenerated, pollen-less and showed lack of cuticle formation. Transverse sections showed normal meiosis till stage 5-6, however, defects in post-meiotic microspore development were found at stage 8-9 in *apa1331*. Measurement of wax and cutin analysis showed a significant reduction in anthers of *apa1331* compared to Wildtype (WT). Quantification of H_2_O_2_ and MDA has indicated the excessive ROS (reactive oxygen species) in *apa1331*. Trypan blue staining, and TUNEL assay revealed cell death and excessive DNA fragmentation in *apa1331.* Map-based cloning and Mutmap analysis identified a candidate gene (*LOC_Os04g40720)* that is a *SUBTILISIN-LIKE SERINE PROTEASE (OsSUBSrP1)* which harbored an SNP (A>G) in *apa1331*. CRISPR-mediated knock-out lines of *OsSUBSrP1* displayed spikelet degeneration comparable to *apa1331*. Global gene expression analysis revealed a significant downregulation of wax and cutin biosynthesis genes e.g., *OsWDA1*, *OsMS2* and *OsCER4* in *apa1331*. Our study reports the novel role of SUBSrP1 in ROS-mediated cell death in panicle development.

**Highlights:**

1. *OsSUBSrP1* plays an important role in maintaining ROS-mediated programmed cell death.
2. *OsSUBSrP1* is essential for apical spikelet development.
3. *OsSUBSrP1* regulates the expression of wax and cutin biosynthesis pathway genes.

## Introduction

Panicle is the reproductive organ in rice which directly determines the grain yield (Zhu *et al*., 2013b). Identification of genotypes with ideal panicle architecture and the responsible genes for this key trait has been the primary focus of rice breeders which significantly contributed in developing ideal plant genotypes with higher grain yield (Huang *et al*., 2009; Huo *et al*., 2017; Wang and Li, 2008). Various genetic and environmental factors regulate the spatiotemporal arrangement of a panicle by affecting its reproductive success and developmental decisions (Teo *et al*., 2014). Apical degeneration of spikelets is highly detrimental to plants and causes a significant reduction in yield (Heng *et al*., 2018; Lu *et al*., 2017; Zafar *et al*., 2020). Although, several genetic factors have been reported that control the apical panicle development in rice. Nevertheless, the molecular basis of mechanisms controlling panicle abortion still needs a comprehensive understanding.

Development of the panicle begins with the transition of the vegetative to the reproductive phase. Various misfortunes at these transitions including shoot, inflorescence and branch meristems could bring severe defects to panicle development (Tanaka *et al*., 2013). *SMALL PANICLE (SP1)* controls the length of the panicle by encoding a nitrate transporter of PEPTIDE TRANSPORTER (PTR) family (Li *et al*., 2009). Similarly, *ERRECT PANICLE 2 (EP), LAX PANICLE1 (LAX1)* and *LAX2* are also found to be involved in the development of axillary and branch meristems formation (Oikawa and Kyozuka, 2009; Tabuchi *et al*., 2011; Zhu *et al*., 2010). *ABERRANT PANICLE ORGANIZATION 1 (APO1)* and *APO2* are reported to work in the temporal regulation of meristems that ultimately affect the inflorescence development (Ikeda *et al*., 2005; Ikedakawakatsu *et al*., 2012). *SQUAMOSA PROMOTER BINDING PROTEIN LIKE 6* (*SPL6)* regulates the signaling outputs by repressing an active transducer INOSITOL-REQUIRING ENZYME 1 (IRE1) of cell death and control panicle development. *SPL6* deficient mutant plants revealed hyperactivation of IRE1 and displayed apical panicle abortion (Wang *et al*., 2018). *ALUMINUM ACTIVATED MALATE TRANSPORTER (OsALMT7)* controls the panicle development by transporting malate to the vascular bundles (Heng *et al*., 2018). A mutation in *CALCINEURIN B-LIKE PROTEIN-INTERACTING PROTEIN KINASE 31 (OsCIPK31)* also displayed apical spikelet abortion phenotype (Peng *et al*., 2018). Mutation in *TUTOU1* that encodes a SUPPRESSOR OF CAMP RECEPTOR (SCAR) like protein also exhibited the apical degeneration phenotype due to disorganization of actin (Bai *et al*., 2015). *OsC6* encodes a LIPID TRANSFER PROTEIN (LTP) and is involved in the post-meiotic development of anthers. Plants silenced for *OsC6* displayed the defective development of pollen exine and tapetum (Zhang *et al*., 2010). *DEGENERATED PANICLE AND PARTIAL STERILITY 1 (DPS1)* played an important role in anther cuticle development, and *dps1* plants showed an accumulation of ROS (reactive oxygen species) in apical spikelets (Zafar *et al*., 2020).

During panicle development, abortion of spikelets frequently occurs either at basal or apical portions of the panicle due to unfavorable conditions (Yao *et al*., 2000). Extreme temperature, malnutrition, and drought stress have been identified as potential factors promoting panicle degeneration (Heng *et al*., 2018; Itoh *et al*., 2005; Smith and Stitt, 2007; Yao *et al*., 2000). Phytohormones regulate the structure of panicle and other plant parts in response to different biotic and abiotic stresses (Alamin *et al*., 2018; Gutierrez *et al*., 2009). Axillary meristems, lateral buds, and reproductive organs are primarily affected by auxin, cytokinin, and various environmental cues (McSteen, 2009). ROS are oxidizing agents and can cause dynamic injury to various biological events (Zhang *et al*., 2013). Impairment in panicle, heading date, plant height and number of grains are associated with ROS accumulation that can cause death to the cellular tissues and machinery (Bai *et al*., 2015; Peng *et al*., 2018; Weng *et al*., 2014). Unbalanced homeostasis of ROS may lead to defects in meiosis and microspore development (Yi *et al*., 2016). Programmed cell death (PCD) is a controlled process and is usually involved in the release and rupture of the nuclear membrane, cytoplasmic shrinkage and swelling of the endoplasmic reticulum (Papini *et al*., 1999).

Anthers are the male reproductive organs inside spikelets which produce pollen grains via meiosis and mitosis which leads to the fertilization of eggs and ultimately seed setting (Ling *et al*., 2015). Anthers are composed of four wall layers; epidermis, endothecium, middle layer, and tapetum. Tapetum is the innermost layer, which serve as a source of nutrients and enzymes during the microspore development (Bedinger, 1992). Normal tapetum development is a key process in the normal pollen development and successful seed setting (Uzair *et al*., 2020). Tapetum development is often associated with a well-developed cuticle, an outer covering that protect anthers from external damages. Cuticle is mainly composed of wax and cutin monomers which consists of several fatty acids. Any abnormality in cuticle development leads to the impaired tapetum ultimately causing the failure in seed set (Zafar *et al*., 2020). Several factors could affect the cuticle development such as high temperature, excessive accumulation of ROS and imbalance of certain hormones (Liao *et al*., 2018; Rang *et al*., 2011; Zhang *et al*., 2017). These factors are either controlled by genetic elements or environmental cues. Wax and cutin are fatty acids and play an essential role in pollen mother cell (PMC) development. C16 and C18 and their derivatives called cutin and constitute the cuticle of anthers (Nawrath, 2006). Anther cuticle and walls share several phenolic and lipidic precursors. Several genes controlling the cuticle and tapetum development have been identified such as *WAX-DEFICIENT ANTHER1 (WDA1)*, *DPS1, POST-MEIOTIC DEFICIENT ANTHER1 (PDA1), ATP BINDING CASSETTE TRANSPORTER G26 (OsABCG26), NO POLLEN1 (NP1), IRREGULAR POLLEN EXINE1 (IPE1), DEFECTIVE POLLEN WALL2 (DPW2),* and *DPW3* have been reported to play their role in anther development by regulating wax and cutin pathway (Chang *et al*., 2016; Chen *et al*., 2017; Jung *et al*., 2006; Mondol *et al*., 2020; Wu *et al*., 2016; Xu *et al*., 2017; Zafar *et al*., 2020; Zhu *et al*., 2013a). *MALE STERILE2 (MS2)* and *DPW2* play their role in developing pollen by regulating the expression of FATTY ACYL TRANSFERASE (Chen *et al*., 2011; Xu *et al*., 2017).

SUBTILISIN-LIKE SERINE PROTEASE (SUBSrP) is a poorly studied family of proteins in plants. However, their role as CASPASES in animals has been widely reported. They display a cleavage specificity and have a particular function in PCD (Vartapetian *et al*., 2011). Some superficial studies have indicated their role as plant-pathogen recognition and determination of silique number (Figueiredo *et al*., 2014; Martinez *et al*., 2015). However, a few plant studies indicated their role in reproductive organs and were also reported as a meiotic protein involved in events of late microsporogenesis (Taylor *et al*., 1997; Yoshida and Kuboyama, 2001).

Current study presents the further genetic analysis and functional validation of our previous preliminary study of a gene mapped on chromosome 4 controlling panicle abortion phenotype (Hou *et al*., 2018). However, the its further follow up genetic analysis and functional analysis was not carried out. Here we present a novel function of *OsSUBSrP1* in panicle development via regulating ROS-mediated cell death. An EMS generated mutant *apa1331,* and CRISPR/Cas9 lines displayed a significant reduction in seed setting rate due to loss of function of *OsSUBSrP1*. Thus, we propose that a mutation in *OsSUBSrP1* triggers the accumulation of ROS and causes cell death in apical spikelets. Our data presents the first insights to the role of *OsSUBSrP1* in seed setting rate by regulating wax and cutin pathway.

## Materials and Methods

### Experimental materials

The panicle mutant *apa1331* was derived from an *indica* maintainer line Yixiang 1B (WT) by EMS (ethyl methane-sulfonate) mutagenesis. *Apa1331* was screened from an M_2_ population, and trait of apical abortion was found stably inherited. In current study, we used *apa1331* mutant that was further subjected to genetic and functional analysis. The *apa1331* mutant was used as female parent, and crossed with the WT of Yixiang 1B (*indica*) and 02428 (*japonica*) cultivars to construct two F2 mapping populations. Plants were grown under natural conditions in the experimental fields at Rice Research Institute, Sichuan Agricultural University, Chengdu (N30.67°, E104.06°), or alternatively at Lingshui (N18.47°, E110.04°), Hainan Province, China.

### Scanning electron microscopy

Scanning electron microscopy was performed as described (Chun et al. 2020). The young panicle buds and shoot apex of the WT and *apa1331* were collected at the primordium differentiation stage. Samples were pre-fixed in pre-cooled 3% glutaraldehyde in 0.1 M phosphate-buffered saline (PBS; 4 mM sodium phosphate, 200 mM NaCl, pH 7.2) at 4°C for overnight. After rinsing with buffer, sample tissues were rinsed again with 5% (w/v) sucrose solution for 5 mints, then dehydrated through a series of ethanol solution (50%, 70%, 85%, 95% and 100%). Samples were then critically dried by a critical point drier, sputter-coated with platinum and observed using a scanning electron microscope (Inspect, FEI, USA).

### DAB staining and quantification of ROS

3,3’-Diaminobenzidine (DAB) staining is used to assess the presence of ROS in the spikelet of WT and *apa1331* by following the previously described method (Wu *et al*., 2017). ROS was quantitatively measured in the form of H_2_O_2_, 100 mg of 6cm fresh panicles of WT and *apa1331* using a ROS assay kit (Beyotime, Shanghai, China). Briefly, panicles were ground into powder form using liquid nitrogen. The powder was further extracted in sodium phosphate buffer at ice for 20 min. The mixture was centrifuged at 12000 g for 15 minutes at 4°C. H_2_O_2_ was quantified from the supernatant measuring the OD value by a spectrophotometer (Thermo Scientific, Varioskan Flash) according to the given manufacturer’s protocol.

### Trypan blue staining

Trypan blue staining is used to the viability of the cells of WT and *apa1331.* Fresh panicles were dipped into the trypan blue solution (2.5 mg/ml trypan blue, 25% lactic acid, 23% phenol and 25% glycerol) and stored at room temperature for 7-12 hours. After that, samples were washed with hot water and de-stained with 50% glycerol for 24 hours. Later, the stained spikelets of WT and *apa1331* were photographed using a microscope (OLYMPUS CX40).

### Quantification of Malondialdehyde (MDA) contents

MDA contents were quantified in 100 mg of 6cm fresh panicles of WT and *apa1331* using an MDA assay kit purchased from Nanjing Jiancheng Bioengineering Institute. The absorbance value of the supernatant was recorded at 450, 532 and 600 nm. MDA quantification was carried out by following a previously described method (Zafar *et al*., 2020).

### Transverse sectioning of anthers

Different stages of anther development were selected from WT and *apa1331* panicles as previously described (Wilson and Zhang, 2009). Paraffin-embedded anthers were cut into 4μ thin slides using a microtome (LEICA RM2255).

### TUNEL assay

TUNEL assay was used to detect cell death and DNA fragmentation as described earlier (Zafar et al. 2020). The samples were selected at the 7 and 12 cm stages of panicle development from WT and *apa1331* and fixed in FAA solution (18:1:1 (v/v) mixture of formalin, acetic acid, and 70% ethanol) for 24 hours. Then, spikelets were dehydrated through a series of ethanol (50%, 70%, 85%, 95%, and 100%) and embedded in paraffin (Roche). Cross-sections of 10 µm thickness were cut using the rotary microtome followed by a dehydration gradient of 95%, 90%, 80%, and 70% ethanol, respectively, 5 min for each. After this, the proteinase K working solution was added and incubated at 37L for 25 min followed by three times washing with PBS (pH 7.4) in a Rocker device (Service Bio TSY-B). The solution was prepared according to the manufacturer’s protocol (Roche, FITC). Then, samples were incubated with DAPI solution at room temperature for 10 min and kept in the dark. DAPI glows blue by UV excitation wavelength 330-380 nm and emission wavelength 420 nm. FITC glows green by excitation wavelength 465-495 nm and emission wavelength 515-555 nm. CY3 glows red by excitation wavelength 510-560 nm and emission wavelength 590 nm. Positive apoptosis cells were green in color and photographed under a confocal laser microscope (OLYMPUS, FV1000)

### RNA extraction and RT-qPCR

A 50-100 mg of fresh tissues from panicle, roots and leaves were taken from WT and *apa1331*. Liquid nitrogen was used as a freezing agent and samples were ground into powder form. After that, different tissues were subjected to TRIZOL (Invitrogen) reagent as per need of experiment and total RNA was extracted and stored at -80 °C.

To reverse transcribe the mRNA for qPCR, RNA was used to make cDNA according to HiScript® IIQ RT SuperMix for qPCR. SYBR Green real-time PCR was used to detect the expression of selected genes in different tissues at different stages. Ubiquitin was used as the internal reference gene, and each sample was set to 3 replicates. Quantitative expression analysis was calculated using 2^-ΔΔ^ method in PCR (Germany) installed with qPCRsoft 3.2 software.

### Genetic analysis of *apa1331*

Map-based cloning was performed with more than 700 SSR markers using 300 mutant individuals selected from F_2_ population. Polymorphic bands were screened from all the chromosomes.

To perform MutMap analysis, F_2_ plants were backcrossed with *apa1331* to generate BC_1_F_2_ population. First, 25 individuals from BC_1_F_2_ population showing the apical degeneration phenotypes were selected for whole-genome resequencing (MutMap). DNA from 25 BC_1_F_2_ plants was pooled and sequenced using the Illumina platform (HiSeq 2500), and WT genomic DNA was also sequenced as a control. The raw data was passed through a set of quality control parameters and only the clean reads were further processed to the next step. The high-quality reads (fastq format) were aligned by BWA (Burrows Wheel Aligner) from each sample against the reference genome (Li and Durbin, 2009). InDels were based on GFF3 files, while ANNOVAR was applied to annotate the SNP (single-nucleotide polymorphisms) in the reference genome (Wang *et al*., 2010). Finally, the delta SNP index was calculated by using the difference of SNP index of pooled samples. PCR products were detected by gel electrophoresis and sequenced (Novogene, Beijing Co., Ltd.). The sequencing alignments and chromatograms of WT and *apa1331* were obtained through the software DNAMAN.

### Development of knock out lines using CRISPR/cas9

To confirm whether *OsSUBSrP1* controls the phenotype of panicle abortion in apa1331 or not, a gRNA sequence (CCACCATGCGCGACCTCAAGTGT) targeted at the first exon of *LOC_Os04g40720* was cloned into pCRISPR/CAS9–MT vector as previously mentioned (Ma *et al*., 2016). After sequence-based confirmation, the vector was transformed into calli of WT. The T_1_ transgenic plants were used for all analysis in this study.

### Transcriptome profiling

Transcriptome analysis was performed using three replicates from the panicles of WT and *apa1331*. Total RNA was extracted from 6cm panicles, when the signs of programmed cell death became visible in the upper panicle of *apa1331* by using the manufacturer’s protocol (mirVana miRNA Isolation Kit, Ambion). A total of 6 samples were sent to OE Biotechnology (Shanghai, China) for complete mRNA transcriptome analysis by following a method reported by (Li and Durbin, 2009). Samples integrity was evaluated using the Agilent 2100 Bioanalyzer (Agilent Technologies, Santa Clara, CA, USA). The samples with RNA integrity number (RIN) ≥ 7 were subjected to the subsequent analysis. According to the manufacturer’s instructions, the libraries were constructed using TruSeq Stranded mRNA LTSample Prep Kit (Illumina, San Diego, CA, USA). Then, these libraries were sequenced on the Illumina sequencing platform (HiSeqTM 2500 or Illumina HiSeq X Ten) and 125bp/150bp paired-end reads were generated.

After getting raw reads, HTSeq was used to analyze the sequences and expressions of all known genes were calculated using the FPKM (fragments per kilobase of transcript per million fragments mapped). All the clusters and DEGs were evaluated by PCA (principal component analysis) and DESeq. All the DEGs were statistically analyzed with cut-off FC >2 and *p*-value ≤0.05. The metric distance, hierarchical clustering, the average linkage was analyzed for DEGs and finally annotated by using GSEA (gene set enrichment analysis). Cytoscape (www.cytoscape.org/) and R-software were used to build the regulatory networks and heat map, respectively. Functional annotation of DEGs was assigned using GO and KEGG pathways terms.

### Analysis of anther wax and cutin

The amount of fatty acids was determined using 25 mg of freeze-dried anthers from WT and *apa1331* as previously described (Liu *et al*., 2021). Briefly, an extraction buffer containing chloroform, methanol and water (8:4:3) was added to freeze the dried powder and vortexed. The mixture was centrifuged at 3000 rpm for 15 minutes. A bottom layer containing the chloroform was transferred to another 10 ml tube. Two ml of solution (KOH-CH_3_OH, 0.5mol/L) was added to the reaction tube and repeated twice. Samples were sealed and oscillated for 1 min in boiling water until the oil droplet disappeared from the surface. After cooling, 2 ml of BF_3_-CH_3_OH for methyl esterification process. 1 ml of n-hexane and 5 ml of saturated NaCl solution were added to the previous mixture, and vail was inserted for fatty acid analysis according to the manufacturer’s protocol (Gas Chromatograph-2010 SHIMADZU).

## Results

### *apa1331* displayed degeneration of apical spikelets

To understand the genetic regulation of panicle development, we isolated a mutant named *apical panicle abortion1331 (apa1331)*, which shows a stable phenotype of panicle degeneration. This mutant was obtained through an ethyl-methanesulfonate (EMS) mutagenized population of an *Indica* rice maintainer line, Yixiang 1B (WT).

At maturity, *apa1331* showed degeneration of apical spikelets and had significantly lower seed yield (Figure 1, A). During reproductive development, the panicle of *apa1331* started to develop the degenerated spikelets at apical portions, while WT did not reveal any degeneration (Figure 1, B). To differentiate the panicle stage, when *apa1331* plants started to show degeneration of apical spikelet. We carefully observed different stages of panicle development (Figure 1C). Until the 5 cm stage of the panicle development, the developmental course of *apa1331* was normal and did not reveal any degeneration signs. After the 6 cm stage of panicle development, degeneration starts to become visible in the apical spikelets of *apa1331*. Panicle began to degenerate apically, even when it was enclosed inside the flag leaf. Panicle degeneration can be seen after the 6 cm stage of panicle development. As the panicle grows in its length, apical spikelets degeneration becomes severer. The ∼6 cm stage of panicle development can be regarded as the point of divergence between *apa1331* and WT. Various agronomic traits, e.g., plant height, number of primary branches, number of tillers, 1000-grain weight did not show any significant differences in *apa1331* compared to WT (Figure 1H-J). However, the number of grains per spike and fertile panicle length was significantly decreased in *apa1331* (Figure 1K-L). Degeneration rate, which is the percentage of degenerated spikelets to the total number of spikelets in a panicle, in *apa1331* was up to 53% (Figure 1M). At the same time, WT did not show degeneration of spikelets. The seed setting rate in *apa1331* was decreased up to 40-53% as compared to WT. Thus, *apa1331* is an apical panicle abortion mutant, which has significantly reduced the number of fertile spikelets that caused a massive loss in grain yield.

**Figure 1.**
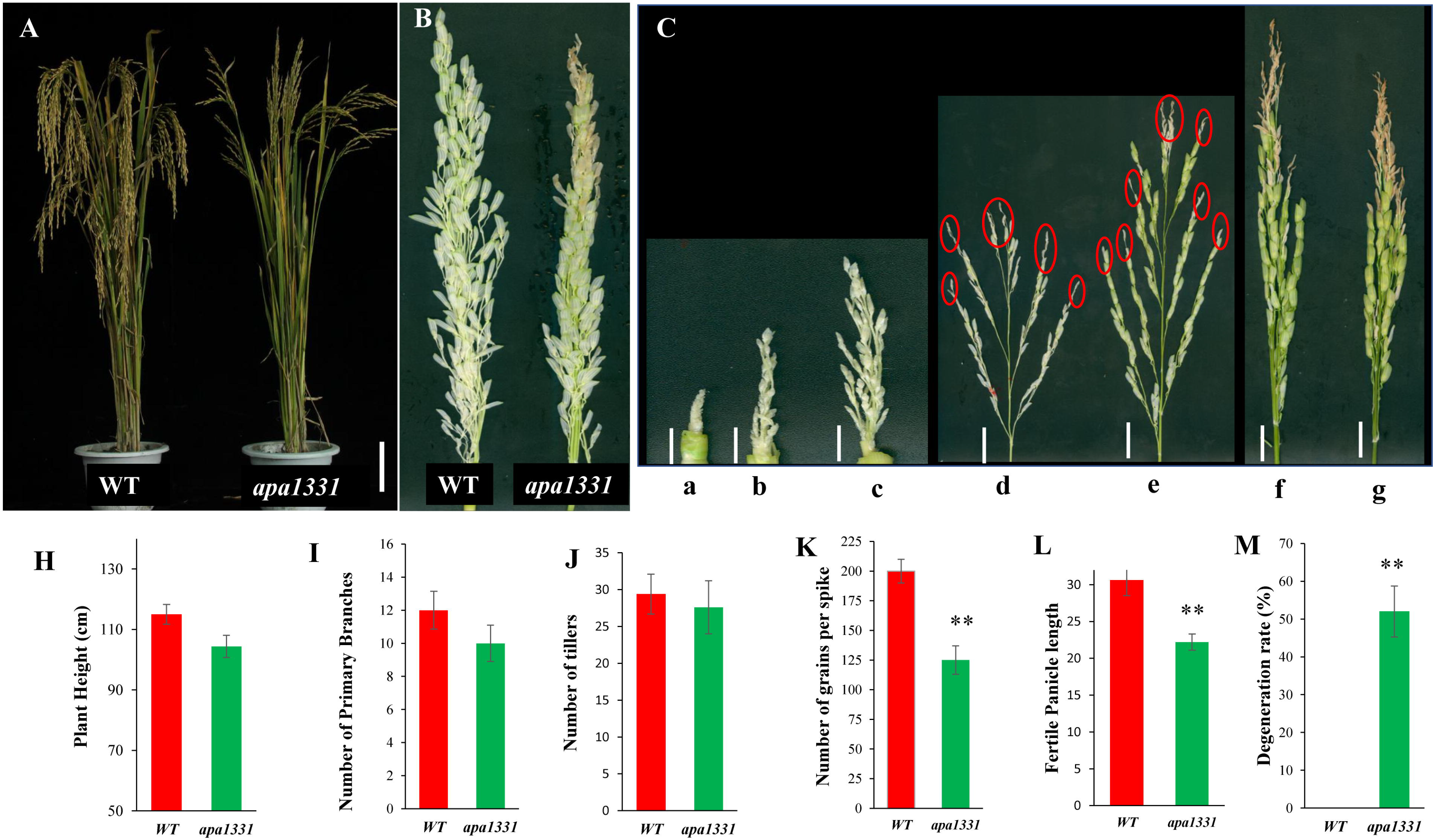
Phenotype and agronomic traits of *apa1331 (apical panicle abortion1331)* **(A)** Plant morphology of a WT plant showing normal panicle development (left) while an *apa1331* shows apical spikelet abortion grown in the field. (right). **(B)** Panicle of WT (left) and *apa1331* (right) before maturity. **(C)** Different stages of panicle of *apa1331* at the length of (a)1 cm, (b) 2.5 cm, (c) 4 cm, (d) 6 cm, (e) 12 cm, (f) 15 cm, and (g) 18 cm. Red circle showing the aborted spikelet. Comparisons of **(H)** plant height, **(I)** number of primary branches, **(J)** number of tillers, **(K)** number of grains per spike, **(L)** fertile panicle length **(M)** and degeneration rate, between WT and *apa1331*. All data are shown in **(H-M)** are denoted, mean±S.D (n=20). Asterisk represents the significant differences calculated by the student’s *t*-test. Where, \**p* <0.05 and \*\**p* <0.01. Bars are equal to 10 cm in **(A)** and 1 cm in **(C).**

### The apical spikelet of *apa1331* are pollen-less and are defective in anther cuticle

To know the basis of the low seed setting rate of *apa1331*, we studied the detailed structure of the apical spikelet using stereo and electron microscopy. The anthers of *apa1331* were degenerated and whitish, while WT anthers were normal and yellowish (Figure 2A-B). WT anthers contain normal, round and viable pollen that were darkly stained in potassium iodide (KI) staining, while *apa1331* spikelets were entirely pollen-less (Figure 2C-D). It is worth mentioning that the basal spikelets of *apa1331* were normal and showed fertile spikelets similar to WT. However, the apical spikelets were sterile. Previous studies have revealed the defective development of anthers cuticle in sterile mutants (Liu *et al*., 2017; Men *et al*., 2017). Accordingly, the scanning electron microscopic (SEM) observation revealed a presence of three-dimensional nano-ridges of cuticle on the surface of WT anther, while that of *apa1331* showed a complete lack of cuticle. The lack of cuticle formation on the surface of *apa1331* anthers has encouraged a look for meiotic development and microsporogenesis (Figure 2E-H). SEM at ∼1.5 cm length showed that *apa1331* did not show any observable aberration at the meristematic level (Figure 2I-J).

**Figure 2.**
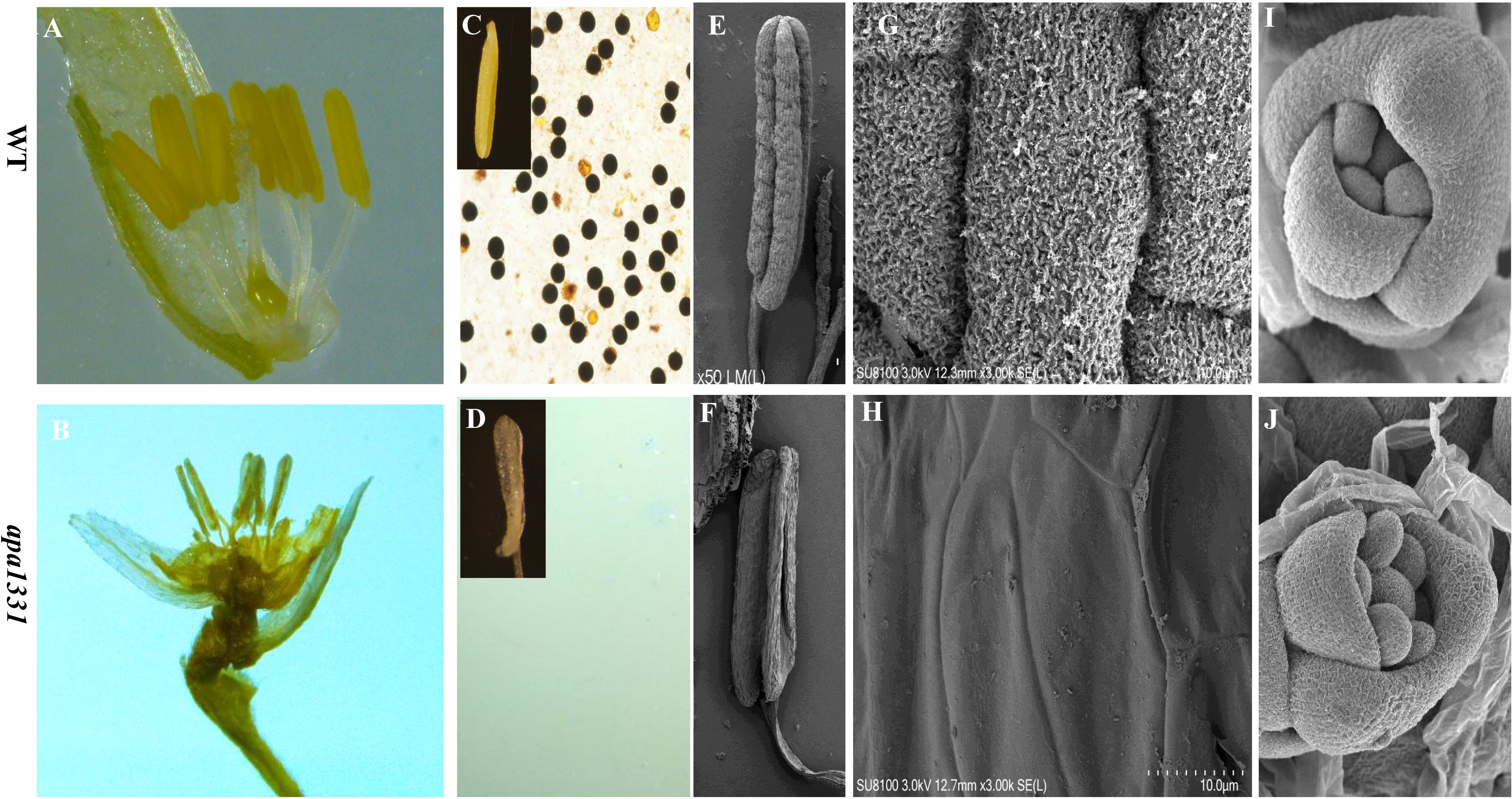
Microscopic examination of Wild type (WT) and *apical panicle abortion1331 (apa1331)* anthers and changes in cuticle structure. **(A-B)** Comparative morphology of WT (up) and *apa1331* (down) flower spikelet obtained by stereomicroscope. **(C-D)** Potassium iodide (KI) staining for pollen viability analysis and its and enlarged view of anther of WT (up) and *apa1331* (down). **(E-F)** Scanning electron microscopy (SEM) morphology of anther of WT (up) and *apa1331* (down). **(G-H)** Closer view of cuticle surface anthers of WT (up) and *apa1331* (down). **(I-J)** Inflorescence meristem development of WT (up) and *apa1331* (down). Bars are equal to 1.5mm in **A-B**, 0.5mm in **C-D,** 50 µm in **G-H** and 500 µm in **I-J.**

### Post-meiotic microspore development was defective in *apa1331*

To reveal cytological defects in the microspore development, cross-sections of different stages of anthers were analyzed as discussed by a previous study (Wilson and Zhang, 2009). Transverse sections of WT and *apa1331* anthers did not reveal any observable difference until stages 6 and 7, both WT and *apa1331* showed the regular structure of all layers e.g., epidermis, endothecium, middle layer, and tapetum (Figure 3A-D). During the early stage 8, the PMCs of WT go under normal meiosis; however, the PMCs nuclei in *apa1331* were not round (asterisk) and were not y callose wall (Figure 3E-F). Moreover, the cytoplasm of *apa1331* was trapped within a boundary. During late stage 8, in *apa1331,* the boundary around the nuclear cytoplasm becomes more evident, and PMC nuclei were irregular (asterisk) and degenerative (Figure 3G-H). In contrast, PMC nuclei of WT were darkly stained and regularly round (Figure 3G). During stage 9, WT tapetum cells were normal, but *apa1331* tapetal cells were enlarged coupled with degenerated microspore cells and nucleus (Figure 3I-J). During stage 10, the tapetum of WT started to degenerate, and microsporocytes were found vacuolated. However, after stage 9, microspore development of *apa1331* was seriously affected and showed severe degeneration of microspores (Figure 3K-L). After stage 9, the microspores of *apa1331* were collapsed and remnants of degenerated microspores were present in the form of cellular debris (Figure 3L). Due to severe degeneration in *apa1331* anthers, we were unable to differentiate later stages of development. The results of cross-sections morphology suggest that microspore development after meiosis was strongly affected in *apa1331* at stage 9.

**Figure 3.**
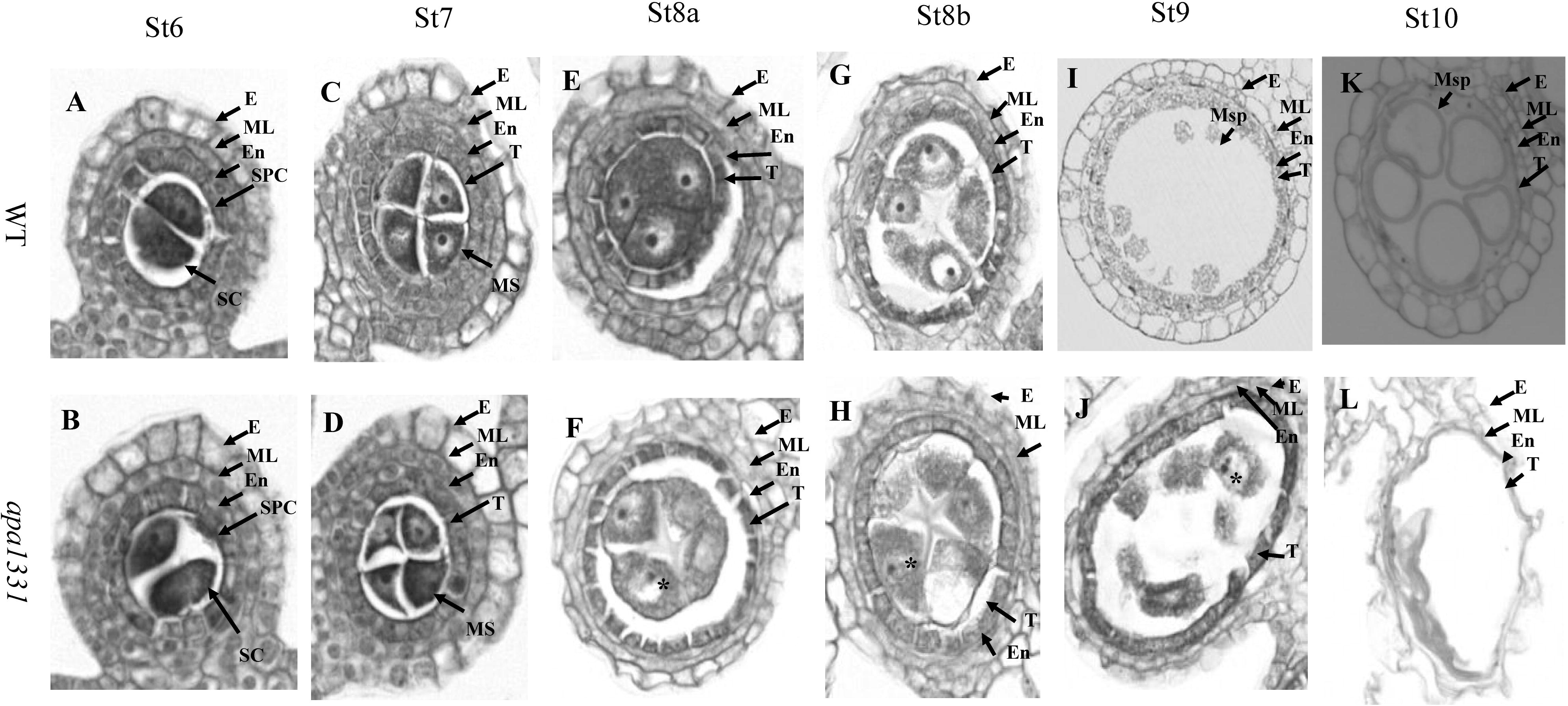
Transverse sections of anthers showing defects microspore development in *apical panicle abortion1331 (apa1331)* **(A-L)** Micrographs of WT and *apa1331* (down) anthers at **(A-B)** St6: showing normal thickening of anthers walls **(C-D)** St7: showing the beginning of defects in *apa1331* development **(E-F)** St8a: showing the trapped cytoplasm in a round enclosure, and degeneration of (pollen mother cell) PMC nucleus in *apa1331*. At the same time, the PMC of WT is regular and round. Asterisk (*) indicates the position of degenerating PMC nucleus **(G-H)** St8b: Trapped cytoplasm and degeneration of PMC nucleus of *apa1331* become more evident at the tetrad stage. **(I-J)** St9: Thickening of T and degeneration of microspore of *apa1331* **(K-L)** St10: At late microspore mother cell development, *apa1331* showed complete degeneration of Msp in the form of degenerated tissue. **E**; epidermis, **ML**: middle layer, **En**; endothecium, **SPC**; secondary parietal cells. **SC**; sporocytes, **T**; tapetum, and **Msp**; microsporocyte. Bars in **A-L** are 25 µm.

### Reduction of wax and cutin contents in *apa1331* anthers

Results of SEM of anthers surface revealed pronounced lack of cutin in *apa1331.* Moreover, the defects in microspore development at the tapetum degeneration stage indicated the difference in fatty acids in *apa1331* and WT. Wax and cutin contents of anthers were measured by gas chromatography, and composition data were presented in the form of percentage as described by a previous method (Liu *et al*., 2021). Results revealed a significant (P<0.01) reduction in total cutin and wax content percentage in *apa1331* as compared to WT (Figure 4A-B). Among saturated fatty acids, palmitic acid (C16:0) and steric acid (C18:0) constituted the cutin monomers and were found significantly (P<0.01) decreased in *apa1331* as compared to WT (Figure 4C). Similarly, the percentage of saturated wax, especially C15:0, C17:0 and C20:0 were significantly reduced in *apa1331* compared to WT (Figure 4D). Among unsaturated cutin C16:1, C18:1nt, C18:1nc and C18:1n3 were significantly decreased in *apa1331* compared to WT (Figure 4E). However, contrary to other fatty acids, unsaturated wax contents e.g., C22:2, C20:5n3, and C22:6n3 were found significantly (P<0.05) increased in *apa1331* as compared to WT (Figure 4F). Consistent with our previous observations, chemical composition analysis of wax and cutin revealed that the candidate gene of *apa1331* plays an essential role in regulating wax and cutin.

**Figure 4.**
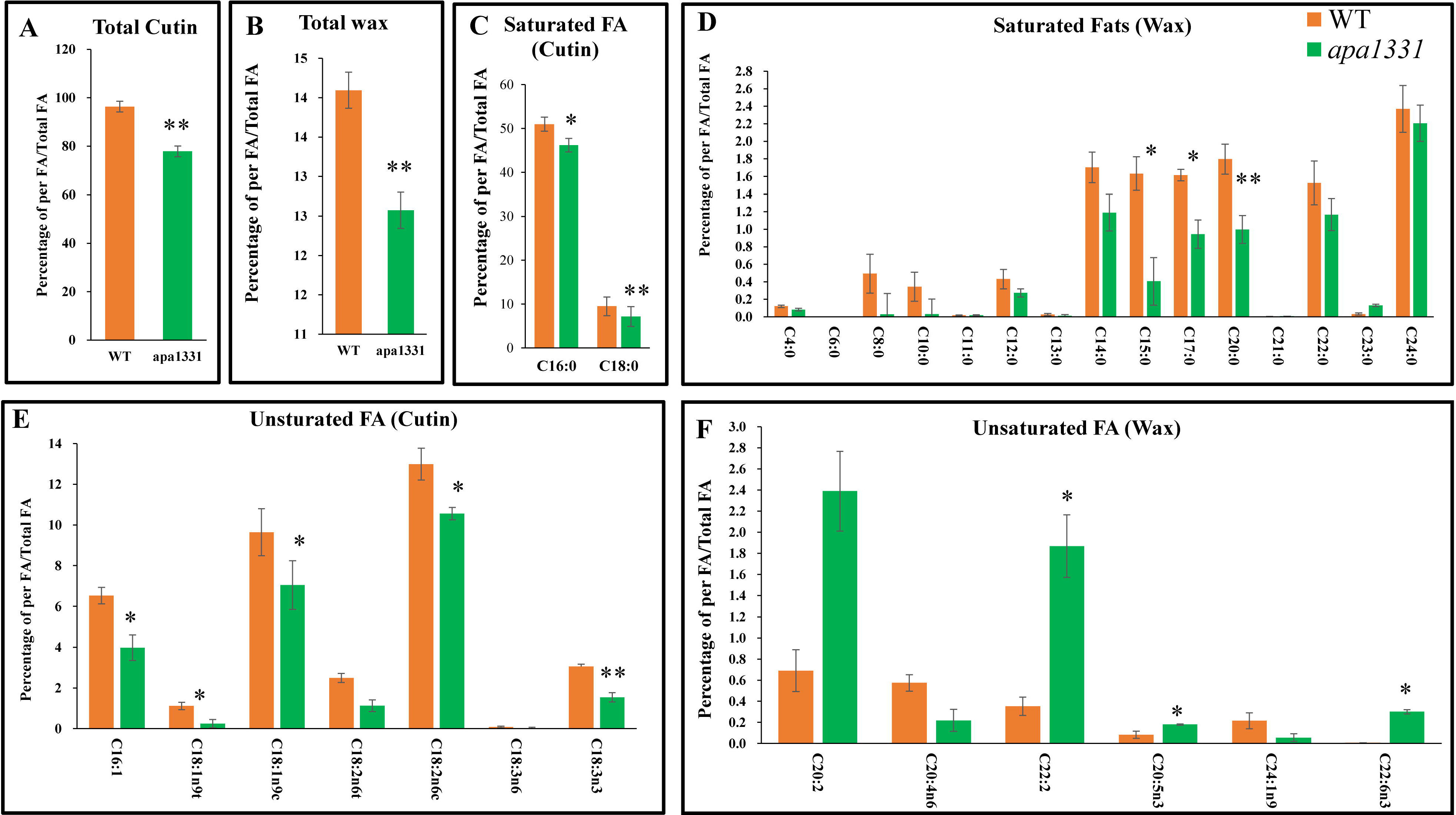
Composition of anther wax and cutin in Wild type (WT) and *apical panicle abortion1331 (apa1331)* **(A)** Total amount of cutin in anthers of WT and *apa1331* **(B)** Total amount of wax in anthers of WT and *apa1331.* **(C)** Amount of saturated fatty acids (cutin) in anthers of WT and *apa1331* **(D)** Amount of saturated fatty acids (wax) in anthers of WT and *apa1331*. (**E**) Amount of unsaturated fatty acids (cutin) in anthers of WT and *apa1331* **(F)** Amount of unsaturated fatty acids (wax) in anthers of WT and *apa1331*. The values represent the average of three individual repeats. Data in **(A-F)** is presented in percentages per fatty acid to total fatty acids. The student’s t-test was used to analyze the significance of data, where \**p* <0.05 and \*\**p* <0.01.

### Apical spikelets of *apa1331* contain a higher level of ROS

Excessive ROS cause damage and induces PCD and are released in plant and animal cells in response to biotic and abiotic stresses. These reactive agents can cause mutilation to the cell wall structure, macromolecules, DNA and proteins (Gomez *et al*., 2003). To further investigate the cause of degeneration of spikelets of *apa1331,* we detected the cellular ROS by DAB staining (Figure 5A). Quantitative measurement of H_2_O_2_ was also significantly (P<0.01) increased in apical spikelet of *apa1331* than WT (Figure 5B). Malondialdehyde (MDA) is an indicator of local ROS production and is involved in the peroxidation of fatty acids (Schmidt and Kunert, 1986). Consistently, quantification of MDA was also significantly increased in *apa1331* spikelets than WT (Figure 5C). Trypan blue staining is used to test the viability of cells (Zhu *et al*., 2016). Dark staining of the apical spikelet of *apa1331* showed cells viability was compromised, while light staining of WT spikelet showed cells were viable (Figure 5D). *CATALASE (OsCATa), OsCATb, OsCATc* are antioxidant enzymes coding genes and involved in the scavenging activities of ROS (Wutipraditkul *et al*., 2011). Relative expressions of *OsCATa, OsCATb,* and *OsCATc* were significantly decreased in *apa1331* spikelets as compared to WT (Figure 5E, Table S1). Together, DAB staining, measurement of H_2_O_2_, relative expression data revealed that excessive accumulation of ROS in *apa1331* caused the degeneration and damage to apical spikelets. At the same time, trypan blue staining has indicated the death of apical spikelets cells.

**Figure 5.**
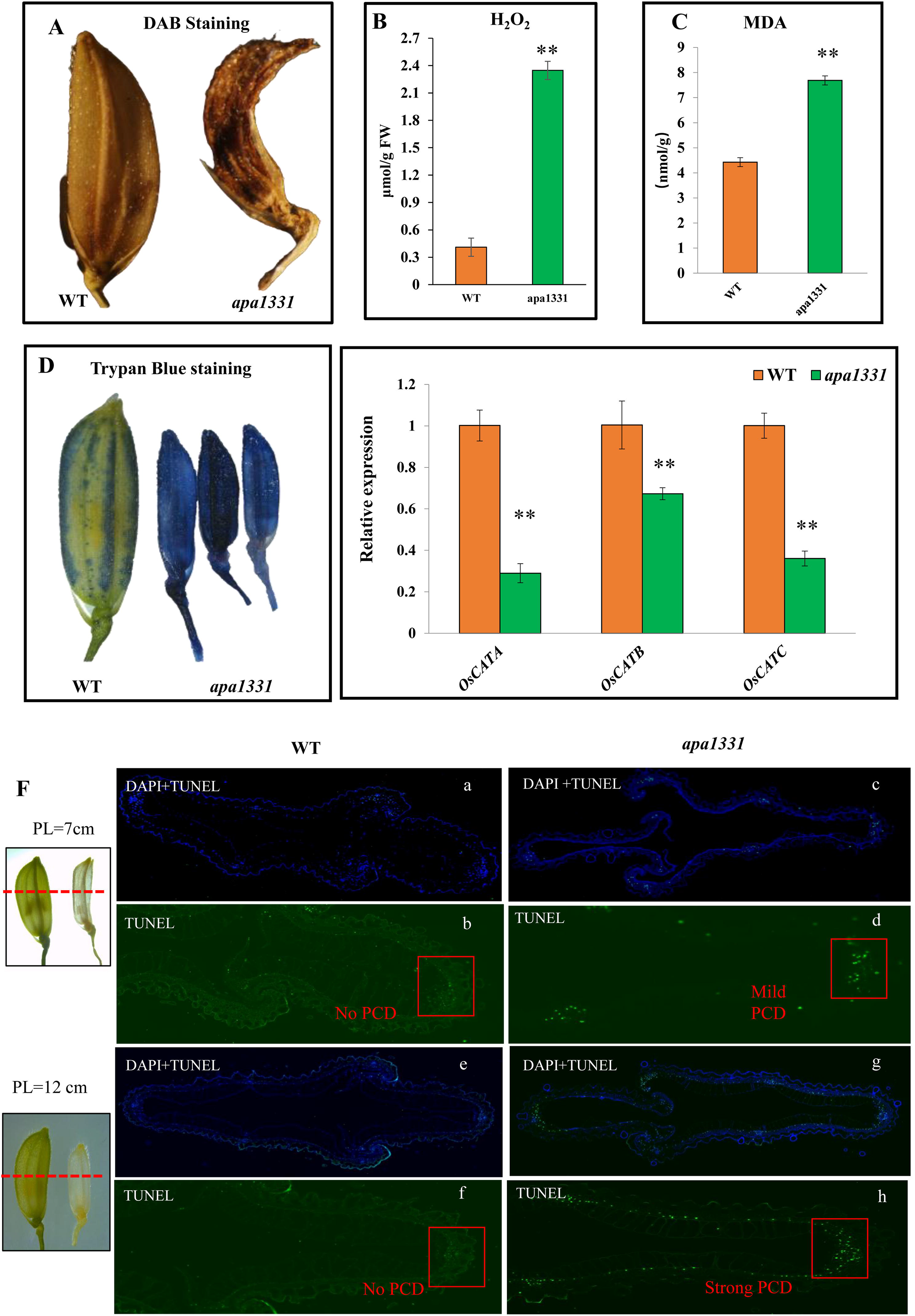
Analysis of reactive oxygen species (ROS), cell death and DNA fragmentation in *apa1331*. **(A)** 3,3-diaminobenzidine (DAB) staining of the spikelet of *apa1331* shows the excessive accumulation of hydrogen peroxide (H_2_O_2_) compared to WT. **(B)** Quantification of H_2_O_2_ was significantly higher compared to WT. **(C)** Quantification of malondialdehyde (MDA) was significantly higher compared to WT compared. **(D)** Trypan blue staining revealed the lower cell viability in apical spikelet of *apa1331* than WT. **(E)** Significant downregulation of *CATALASES,* e.g., *OsCATa, OsCATb* and *OsCATc* in the panicle of *apa1331* than WT. **(F)** TUNEL (terminal deoxynucleotidyl transferase dUTP nick end labeling) assay revealed mild DNA fragmentation at panicle length of 7cm stage. a,c,e and g showed the merged view of DAPI (blue) and (FITC-TUNEL) fluorescein iso-thiocyanate (green dots, displaying the positive signals of apoptosis). TUNEL of 7 and 12 cm panicle from WT did not show any positive signal showing the absence of programmed cell death and DNA fragmentation (b and f). **(E and F)** TUNEL assay in *apa1331* panicle at 7 cm. The presence of weak signals of FTIC (green dots) at the 7 cm stage (d) and strong signals at the 12 cm stage (h) showed the excessive PCD and DNA fragmentation. **(G)** TUNEL assay at 12 cm showed the intensive signals of FITC displaying severe DNA fragmentation and programmed cell death in *apa1331.* The red square line shows the signal of TUNEL.

### Apical spikelets of *apa1331* showed increased cell death

Trypan blue staining has indicated the cell death of the apical spikelets of *apa1331*. We hypothesized that the apical spikelets of *apa1331* would have abnormal PCD and DNA fragmentation. To test this hypothesis, we performed a TUNEL (terminal deoxynucleotidyl transferase-mediated dUTP nick-end labeling) assay to detect apoptosis in spikelets by following a previous method (Zafar et al. 2020). Panicles of 7 and 12 cm length from WT and *apa1331* were used for the TUNEL assay (Figure 5F). Panicles at both 7 and 12 cm stages showed a minor or no positive TUNEL signal, which revealed the absence of abnormal PCD in WT (Figure 5F, b and f). However, the apical spikelet of 7 cm panicle of *apa1331* showed a weak positive signal of TUNEL, revealing the start of mild PCD (Figure 5F, d). Apical spikelet of 12 cm panicle of *apa1331* showed the vigorous-intensity of positive TUNEL signal indicating the DNA fragmentation and vigorous PCD (Figure 5f, g). These findings suggest that enhanced PCD and DNA fragmentation has lead to degeneration of apical spikelet of *apa1331*.

### Candidate gene analysis of *apa1331*

To find the candidate gene of the characterized mutant phenotype, we developed two F_2_ mapping populations. These mapping populations were derived by crossing *apa1331* with an *indica* cv. Yixiang 1B and a *japonica* cv. 02428, respectively. In F_1_ generations of both populations, all plants did not show degeneration phenotype. Genetic analysis of these population revealed that a single recessive gene controls the phenotype of panicle degeneration in *apa1331(Hou et al., 2018)*.

Primary mapping of *apa1331* revealed that the candidate gene resides between the SSR marker 4-9 and 4-10 on the short arm of chromosome 4 (Figure 6A). The gene was narrowed down between region of RM252 and RM5979 markers in a distance of 16.4cM (Hou *et al*., 2018). Due to the further unavailability of InDel markers in this region, we performed MutMap assay to find any mutation in the primary mapped region (Abe *et al*., 2012). MutMap whole genome sequencing (Illumina HiSeq 2500 platform) revealed an SNP index of 1 due to the presence of an SNP in *LOC_Os04g40720 (Hou et al., 2018).* Consistent with the primary mapping by SSR markers, Mutmap results showed an SNP (A > G) was located between the previously mapped region between RM252 and RM5979. According to MutMap, the candidate gene of *apa1331* phenotype was *LOC_Os04g40720*; it has six exons and present in reverse order in the MSU database. The mutation was located in the 4^th^ exon of *LOC_Os04g40720,* changing the 2476^th^ codon from AAC to GAC (Figure 6B). This SNP has ultimately changed the amino acid asparagine into aspartic acid. According to Rice Genome Annotation Project (http://rice.plantbiology.msu.edu/), *LOC_Os04g40720* is annotated for a *SUBTILISIN LIKE SERINE PROTEASE (SUBSrP),* and it contains 1393 amino acids (Ouyang *et al*., 2007). SUBSrP further consisted of a conserved CC (coiled-coil) domain of 21 amino acids, from 455-475 (Figure 6C). To further verify the MutMap candidate gene analysis, we cloned the candidate gene ’*LOC_Os04g40720’* in *apa1331* and WT and sequenced it. The comparative chromatogram of WT and *apa1331* validated the presence of an SNP as revealed by MutMap analysis (Figure 6D). Hence, we tentatively named the protein SUBSrP1 and its gene as Os*SUBSrP1*.

**Figure 6.**
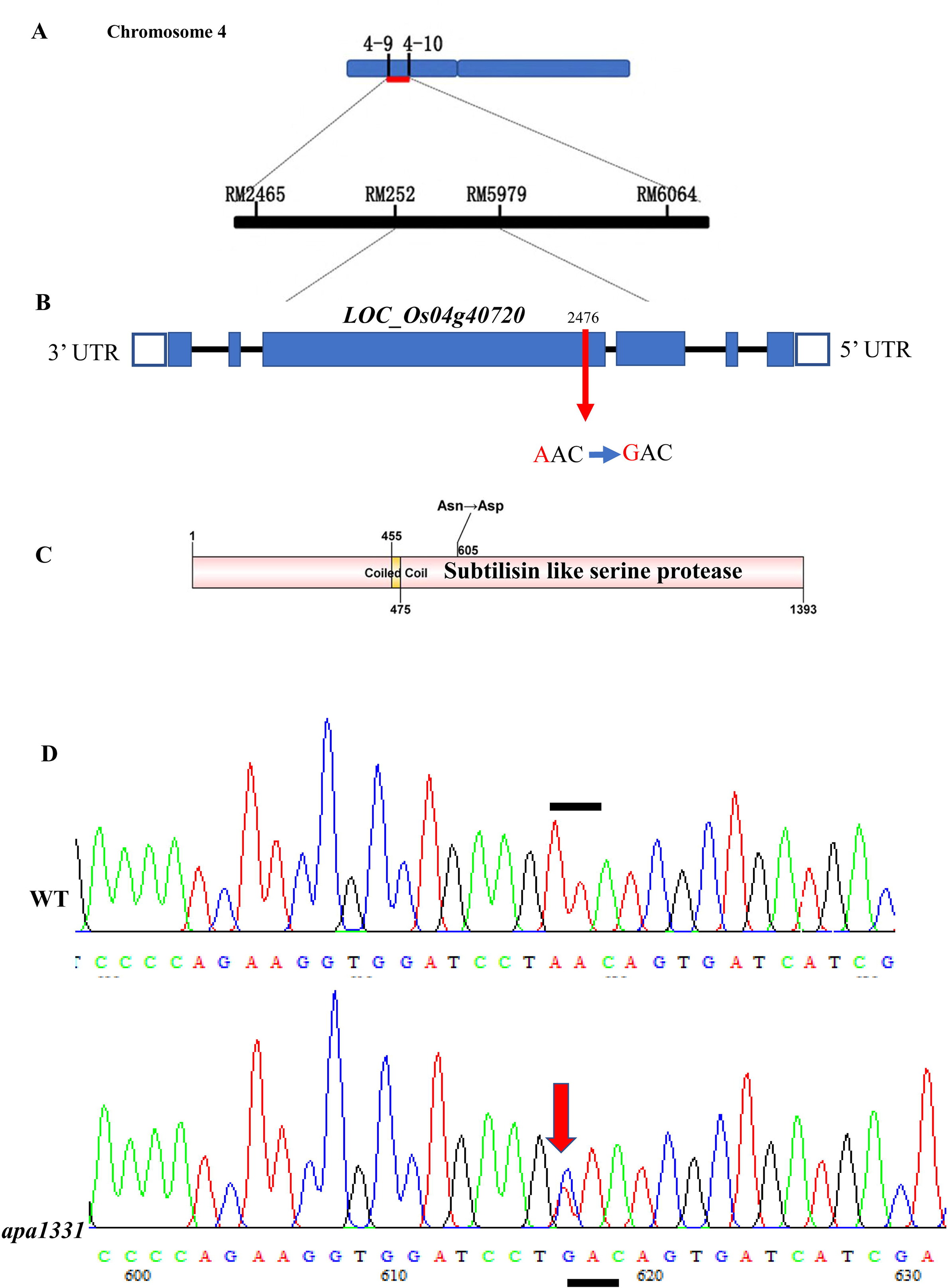
Fine mapping and genetic analysis of *OsSUBSrP1 (SUBTILISIN-LIKE SERINE PROTEASE 1)* **(A)** Primary mapping of the *OsSUBSrP1* controlling *apa1331* phenotype on chromosome 4 between SSR markers 4-9 and 4-10 **(B**) Genomic structure of *OsSUBSrP1 (LOC_Os04g40720)* and position of SNP, where blue boxes are representing exons, blue lined boxes are up and downstream sequences, while black lines are indicating introns. The red arrow shows the position of SNP that was harbored in the 4^th^ exon of *OsSUBSrP1* **(C)** SUBSrP1 consists of 1393 amino acids, and an SNP caused the non-synonymous substitution (Asn > Asp) of 605^th^ amino acid. Coiled-coil domain (CCD) ranged from 455 to 475^th^ position of amino acid. **(D)** shows the comparative chromatograms, cloned and sequenced from WT (up) and *apa1331* (down), where the red arrow shows substituted amino acid, and the black dotted above the WT chromatogram sequence.

These results showed that *OsSUBSrP1* is a recessive gene and controlling the phenotype of apical spikelets of *apa1331*.

### CRISPR-*Cas9* targeted knock outlines showed the phenotype of apical spikelet abortion

To further validate the function of *OsSUBSrP1*, we transformed a CRISPR-*Cas9* PAM sequence (CCACCATGCGCGACCTCAAGTGT) of sgRNA into WT, targeted at the first exon of Os*SUBSrP1* (Figure 7A). Three independent Knock out (KO) lines showed the degeneration of apical spikelets in T_2_ generation. Sequencing revealed deletion of a one bp (G), one bp substitution (C to T), and one bp substitution (G to A) in KO-1, KO-2, and KO-3, respectively. KO-1, KO-2 and KO-3 showed the degeneration of apical spikelets, while the WT did not show any degeneration phenotype at the heading stage (Figure 7B). The relative expression of *OsSUBSrP1* in KO-1, KO-2, and KO-3 was significantly (P<0.01) down-regulated compared to WT. Similar to *apa1331*, the number of seeds per panicle and fertile panicle length was significantly (P<0.01) decreased in KO-1, KO-2, and KO-3 compared to WT. Overall, genetic analysis of *apa1331* and CRISPR/Cas9-targeted mutagenesis of KO lines revealed that mutation in *OsSUBSrP1* causes a significant decrease in seed setting and fertile panicle length.

**Figure 7.**
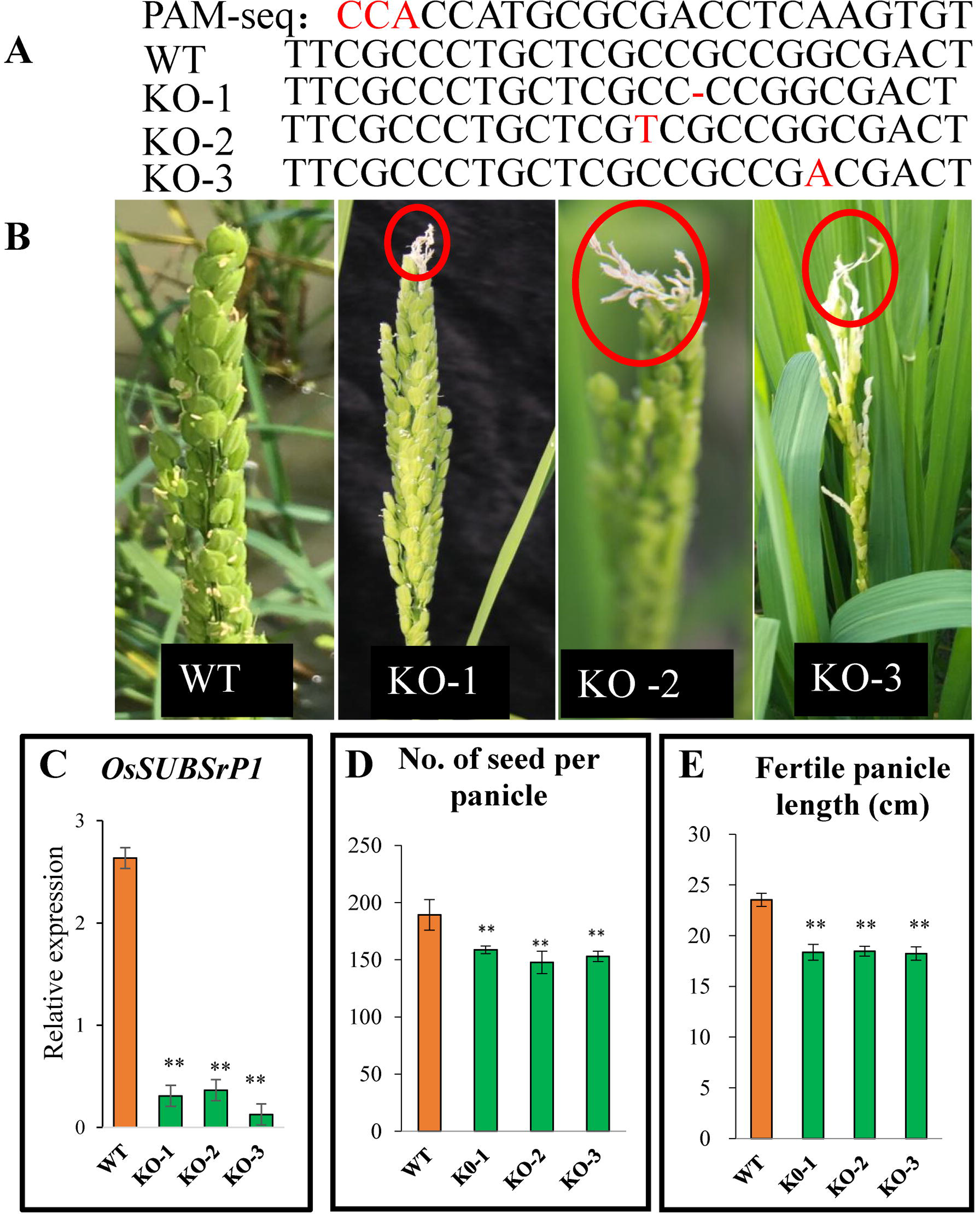
CRISPR/Cas9 mediated knock out lines of *OsSUBSrP1* showed the apical spikelet degeneration. **(A)** The sequence of Photospacer adjacent motif (PAM), Wild type (WT), Knock-out line (KO-1), KO-2, and KO-3. Where red color (-) showed deletion of one bp, C to T, G to A in KO-1, KO-2, and KO-3, respectively. **(B)** Spikelet of KO-1, KO-2, and KO-3 showing apical abortion, while the WT did not show any degeneration phenotype. **(C)** Relative expression of *OsSUBSrP1* was significantly (P<0.01) downregulated in KO-1, KO-2, and KO-3. **(D)** Number of seeds per panicle was significantly ((P<0.01)) decreased in KO-1, KO-2, and KO-3. **(E)** Fertile length of panicle was significantly decreased due to apical panicle degeneration in KO-1, KO-2, and KO-3. The student’s t-test was used to analyze the significance of data, where \*\**p* <0.01.

### Global gene expression analysis revealed *OsSUBSrP1* mediated downregulation of wax and cutin biosynthesis pathway genes

Abnormal PCD and elevated ROS levels can affect the expression of many wax and cutin pathway genes (Zafar *et al*., 2020). To elucidate the transcriptional changes in functional pathways due to elevated ROS levels caused by a mutation in *OsSUBSrP1*, a comparative transcriptome profiling among the WT and *apa1331* panicles was performed using RNA sequencing Illumina Platform (HiSeqTM 2500). Three individual repeats were selected from WT and *apa1331* panicles at the 6 cm stage (point of divergence between WT and *apa1331*). High-throughput RNA sequencing generated high-quality data with an adequate sequencing depth. PCA (principal component analysis) revealed 76% and 19% variance between WT and *apa1331* samples at PC1 and PC2, respectively (Figure 8A). The heat map showed 3168 differentially expressed genes (DEGs) between *apa1331* and WT (Figure 8B). Among 3168 DEGs, 1935 and 1233 DEGs were found down and up-regulated in *apa1331* compared to WT, respectively (Figure 8B). We further subjected the significantly [fold change (FC) > 2 and a *p*-value less than 0.05] DEGs to GO (gene ontology) and KEGG (Kyoto encyclopedia of genes and genomes) pathway functional classification.

**Figure 8.**
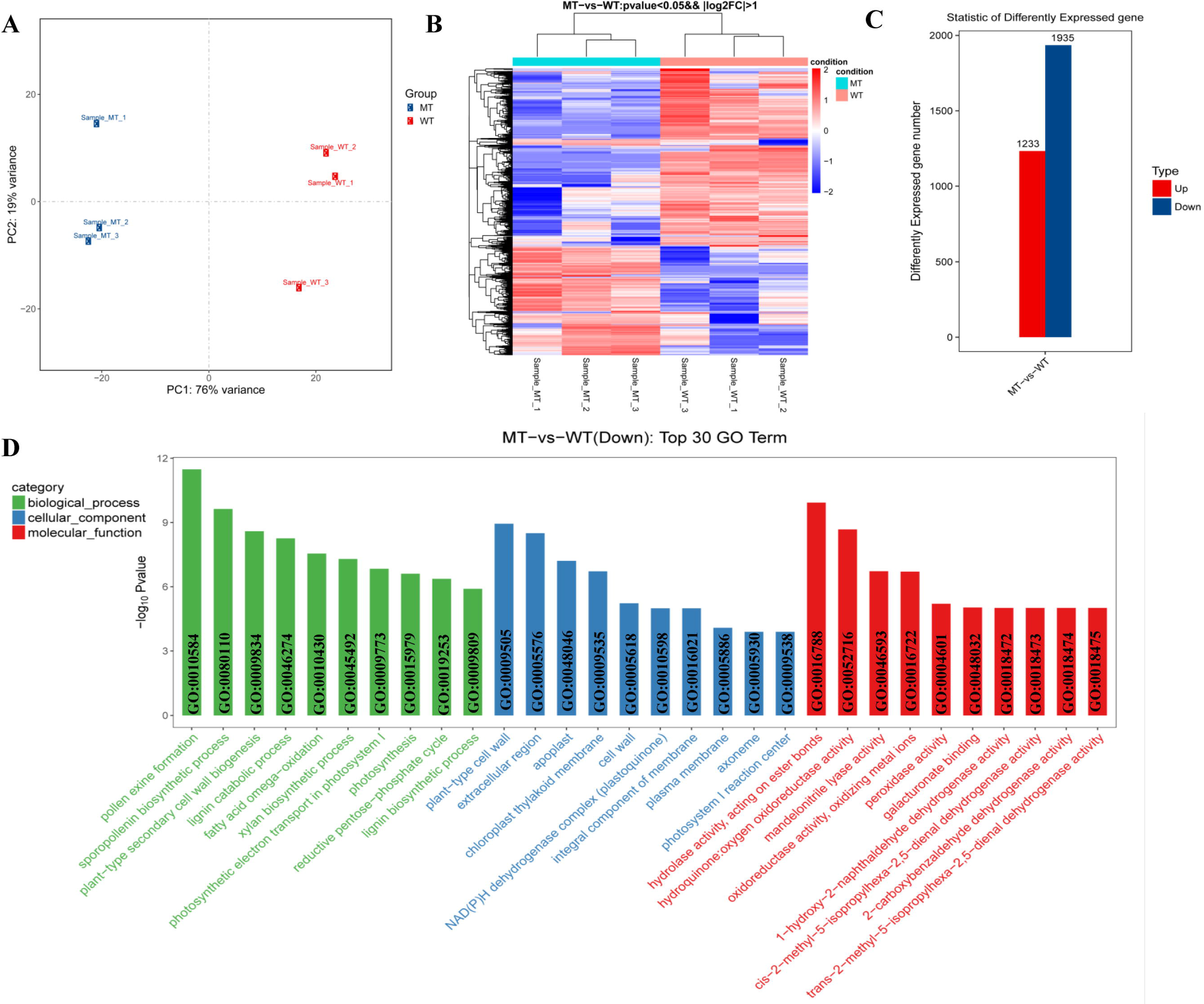
Global gene expression analysis of Wild type (WT) and *apical panicle abortion1331 (apa1331)* **(A)** Principle coefficient analysis (PCA) shows 76% and 19% variations between (PC1) and within (PC2) the samples, respectively. **(B)** Heat map showing the differentially expressed genes (DEGs) with log2FC >1 at *p*-value <0.05 in *apa1331* compared to WT **(C**) Graph showing statistics of DEGs between WT and *apa1331,* where red and blue color are showing the number of up and down-regulated genes, respectively. **(D)** Comparison of top 30 downregulated GO (gene ontology) terms enrichment analysis. The x-axis shows the GO terms of biological processes, cellular components and molecular function in green, blue and red color, respectively, while the y-axis shows the -log_10_ *p*-value of differentially expressed GO terms between WT and *apa1331*.

Further, GO classification of these significantly different DEGs between *apa1331* and WT was categorized into biological processes, cellular components and molecular functions. Among the top 30 down-regulated DEGs, pollen exine function (GO:0010584) was the most significantly down-regulated biological process in *apa1331* (Figure 8D). While other important biological processes associated with these DEGs were enriched in the sporopollenin biosynthetic process (GO:0080110), fatty acid omega oxidation (GO:0010430), and lignin biosynthetic process (GO:000980). While most significantly downregulated cellular components and molecular functions associated with these DEGs were ’plant-type cell wall (GO:0009505)’ and ’hydrolase activity, acting on ester bonds (GO:00016788), respectively. GO classification indicates the downregulation of pollen exine, sporopollenin, fatty acid omega oxidation related biological processes in *apa1331* panicle.

We mapped all DEGs to their respective KEGG classification to get further insights into the specific pathway that regulates these biological processes. Among them, cutin, suberin, and wax biosynthesis pathway (K00073) was the most relevant KEGG pathway with the highest (15.3%) number of DEGs (Figure 9). 13 genes from 3168 DEGs were involved in the wax and cutin biosynthesis pathway of fatty acids. Among the 13 genes, 12 genes *(Os11g0507200, Os01g0854800, Os06g0254700, Os06g0254300, Os10g0486100, Os08g0401500, Os04g0512200, OsWDA1, OsMS2 (MALE STERILITY2), OsCER4, Os07g0416100* and *Os08g0298700)* were found significantly downregulated. While only one gene of *Os02g0814200,* annotated for aldehyde decarbonylase (EC: 4.1.99.5) was found significantly up-regulated (FC=3.341) in MT as compared to WT. The heat map of these 12 downregulated and one upregulated gene represents the significant changes in their expression in *apa1331* compared to WT. Among the down-regulated genes e.g., *Os11g0507200* encodes a transferase family protein and is involved in the biosynthesis of unsaturated fatty acids, especially cutin and suberin. *Os01g0854800* and *Os10g0486100* encodes a cytochrome P450 protein and are involved in the long-chain fatty acid mono-oxygenase activity (EC:1.14.14.80). *Os06g0254300* encodes a peroxygenase and is involved in the cutin, suberin and wax biosynthesis pathway. *OsWDA1, OsMS2,* and *OsCER4* have been reported to regulate the wax and cutin biosynthesis and pollen development in rice (Jung *et al*., 2006; Shi *et al*., 2011; Wang *et al*., 2017).

**Figure 9.**
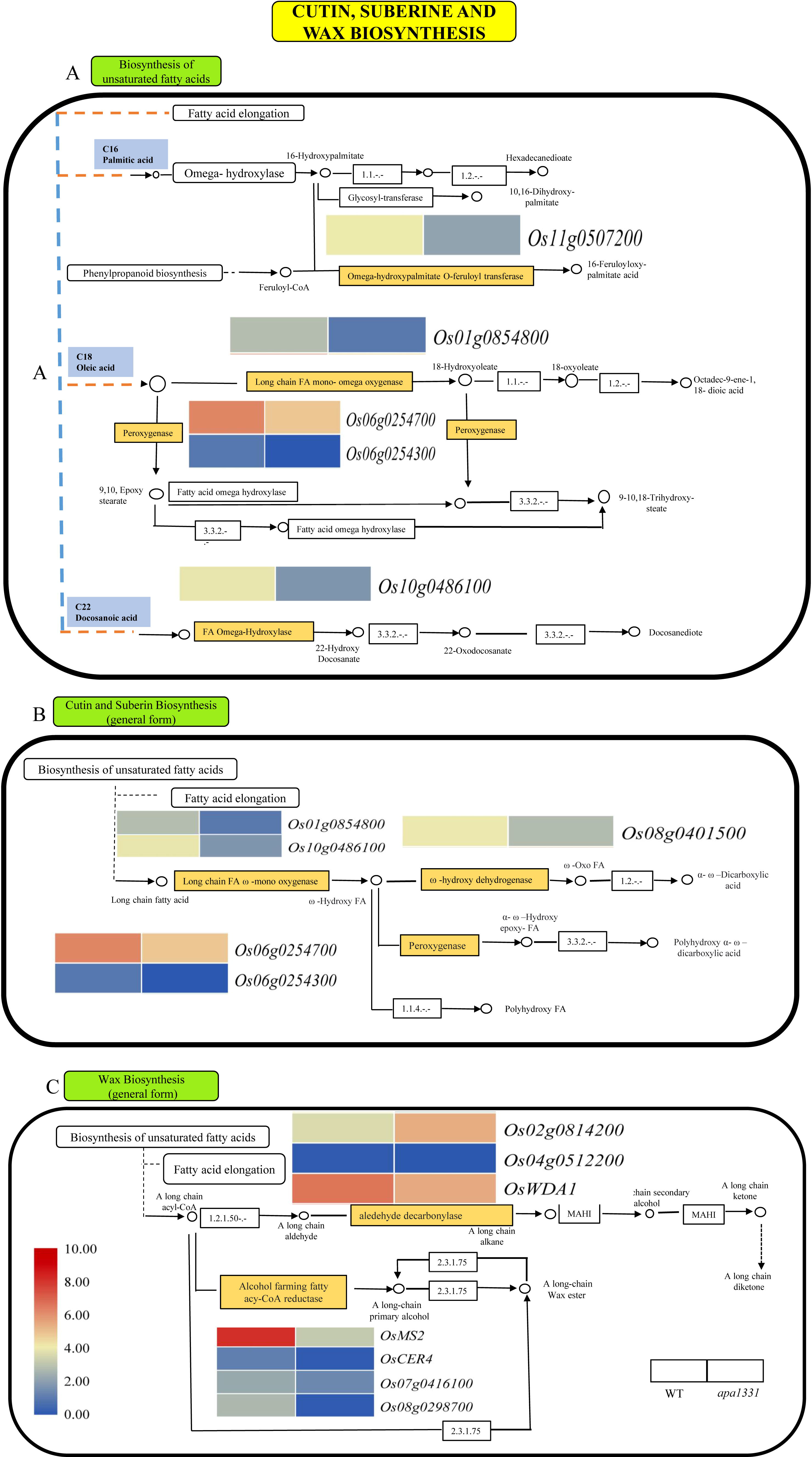
*SUBTILISIN-LIKE SERINE PROTEASE 1 (OsSUBSrP1)* regulates the expression of wax, suberin, and cutin biosynthesis pathway genes. **(A)** Comparative transcriptome analysis of Wild type (WT) and *apical panicle abortion1331 (apa1331)* revealed the downregulation of genes involved in the biosynthesis of the unsaturated fatty acids. **(B)** Comparative transcriptome analysis of WT and *apa1331* revealed the downregulation of genes involved in the biosynthesis of cutin and suberin biosynthesis. **(A)** Comparative transcriptome analysis of WT and *apa1331* revealed the up and downregulation of genes involved in the wax biosynthesis. The heat maps showed the normalized expression value of fragments per kilobase of transcript per million (FPKM) of given genes. Rectangles filled with brown color show the downregulation in **A, B** and **C**. Legend shows the normalized expression value of the presented gene (left to a heat map), where dark red shows the highest and dark blue shows the lowest expression in *apa1331* compared to WT.

To validate the expression of DEGs obtained in RNA-seq data, we randomly selected five genes among up and down-regulated DEGs and determined their relative expression by RT-qPCR (Figure 1S, Table 1S-2S). Consistently, qRT-PCR results were consistent with RNA-seq data.

These data revealed the mutation in *OsSUBSrP1* causes the significant downregulation of wax and cutin biosynthesis genes that lead to the degeneration of apical spikelets in *apa1331*.

## Discussion

The panicle is an essential organ for the reproductive yield of a plant. Panicle abortion is a serious physiological defect and causes heavy losses to the grain yield (Heng *et al*., 2018). Apical spikelet degeneration has been reported to cause a significant reduction in seed setting rate (Peng *et al*., 2018; Zafar *et al*., 2020). Recently, several mutants, e.g., *dps1, rbh1 (rice bald head 1), totou1,* and *spl6* showed the phenotype of apical spikelet degeneration (He *et al*., 2021; Wang *et al*., 2018; Zafar *et al*., 2020). None of the previously reported mutants encode a SUBSrP for apical spikelet development. In the present study, we characterized a novel gene *OsSUBSrP1* that encodes a SUBSrP1. It has been widely reported to control cell death events in animal cells. However, its function has not been reported in plants so far. An SNP in *OsSUBSrP1* caused the dysregulation of SUBSrP1 and produced degeneration of apical spikelets. Thus, we characterize the role of *OsSUBSrP1* in apical spikelet development for the first time.

### *OsSUBP1* is involved in anther cuticle formation and regulate the wax and cutin biosynthesis pathway

Fatty acids are essential molecules required during the degeneration of anther tapetum (Liu *et al*., 2017). Anthers of apical spikelet of *apa1331* were pollen-less and sterile and their SEM revealed significant defects in cuticle formation (Figure 2E-G). Transverse sections of the apical spikelet of *apa1331* revealed the abnormal thickening of tapetum and degeneration of PMC nucleus at stage 9 (Figure 3K-L). Previous studies showing severe abnormal developments at later meiosis stages also led to abnormal pollen development (Liu *et al*., 2017; Yi *et al*., 2016). Anther cuticle is composed of cutin and wax, which are long-chain fatty acids, alcohols and alkanes (Samuels *et al*., 2008). Moreover, comparative fatty acid profiling revealed a significant decrease in saturated and unsaturated wax and cutin in the apical spikelet of *apa1331* compared to its WT (Figure 4). Previous studies revealed that the decrease of wax and cutin contents could cause defects in pollen development (Chen *et al*., 2011; Liu *et al*., 2017; Men *et al*., 2017; Zhang *et al*., 2010). A significant decrease in wax and cutin contents also suggests the substantial loss of water that has caused the apical spikelet degeneration in *apa1331* (Aharoni *et al*., 2004). Transcriptome profiling of apical spikelets revealed the significant downregulation of wax and cutin pathway genes in *apa1331* compared to WT. Downregulation of wax and cutin genes supports the observation of decreased anther cuticle formation (Figure 9). Global gene expression analysis revealed that 13 genes were directly involved in the metabolism of fatty acids. *OsWDA1, OsMS2* and *OsCER4* have already been reported to regulate wax biosynthesis, pollen exine and anther walls formation (Chen *et al*., 2011; Jung *et al*., 2006; Liu *et al*., 2021; Wang *et al*., 2017).

*Os01g0854800, Os06g0254300* and *Os10g0486100* also encode different enzymes involved in regulating cutin, suberin, and wax biosynthesis pathway. A considerable decrease in saturated and unsaturated wax and cutin in *apa1331* was likely due to downregulated genes mentioned above. These findings support that *OsSUBSrP1* is essential for normal development of PMC and wax cutin pathway. Loss of function of *OsSUBSrP1* causes the improper development of wax and cutin that ultimate lead to the production of aborted apical spikelet in *apa1331.* Our data suggest *OsSUBSrP1* plays an essential role in the regulation of anther cuticle formation by regulating the expression of wax and cutin biosynthesis genes.

### *OsSUBSrP1* regulates ROS-mediated cell death

DAB staining and measurement of H_2_O_2_ revealed the excessive accumulation of ROS in apical spikelets of *apa1331* (Figure 5). Enhanced level of MDA and decreased relative expression level of *OsCATa, OsCATb* and *OsCATc* revealed the excessive peroxidation of fatty acids and ROS imbalance in *apa1331* (Figure 5E). Dark staining of trypan blue and strong positive TUNEL signals revealed DNA fragmentation and cell death in *apa1331*(Figure 5F). Previous studies have reported that the excessive occurrence of ROS causes cell death in the apical spikelet (Heng *et al*., 2018; Peng *et al*., 2018). Enhanced ROS levels and increased PCD in the apical spikelet indicate their correlation with the apical spikelet abortion. Excessive PCD have been reported to play its role in panicle development and spikelet abortion (Jung *et al*., 2006; Zhang *et al*., 2010). Enhanced PCD is usually accompanied due to disturbance in the ROS homeostasis or external stimuli (Mittler *et al*., 1997). Aborted pollen grains, failure of normal microspore development and abnormal anther development have been reported due to excessive accumulation of ROS (Hu *et al*., 2011; Zafar *et al*., 2020). ROS are important signaling molecules involved in the rupture of tapetum for pollen grain development but its accumulation beyond the threshold would produce an oxidative stress to the important cellular components (Duan *et al*., 2014). Hence, it is logical to refer that apical sterility in *apa1331* was caused due to excessive burst of ROS and abnormal cell death occurrence. Our findings provide the basis of the molecular mechanism in which homeostasis in ROS-mediated PCD was possibly dysregulated due to the mutation in *OsSUBSrP1*. Our data support that apical spikelet degeneration onset was started due to mutation of *OsSUBSrP1* and validate the presence of potential correlation of cell death and ROS homeostasis for panicle development.

### *OsSUBSrP1* encodes a SUBSrP1 and is essential for normal panicle development

Genetic analysis of *apa1331* has revealed that it bears an SNP (A>G) in the 4^th^ exon that causes the dysregulation in its candidate gene *OsSUBSrP1* (Figure 6)*. OsSUBSrP1* encodes a SUBSrP1, and its role in the plants is poorly studied. A study has indicated the secretion of a serine proteinase at late microsporogenesis and primary expression in tapetal cells (Taylor *et al*., 1997). This study has indicated the sequence of serine proteinase shows that it is a preproprotein, which acquired the glycans during moving from Golgi body and mitochondria to other organelles. A study revealed the preferential expression of a preproprotein *RICE SUBTILISIN-LIKE SERINE PROTEASE 1* (*RSP1)* in ovaries and pistils (Yoshida and Kuboyama, 2001). Few family members have also been reported to be differentially expressed in tomatoes (Jordá *et al*., 1999). *SENESCENCE-ASSOCIATED SUBTILISIN LIKE PROTEASE (SASP)* reported in Arabidopsis and its loss of function mutant showed increased inflorescences branches at the reproductive stage (Martinez *et al*., 2015). This study has also reported that subtilisin-like *SASP* function is conserved at least between rice and Arabidopsis. However, functional characterization of any individual plant serine protease has not been reported so far. Nonetheless, these findings have indicated the presence of SUBSrP in rice and preferential expression in flower development (Martinez *et al*., 2015; Yoshida and Kuboyama, 2001). Our study presents that the loss of function of *OsSUBSrP1* produced apical spikelet degeneration. Our study will to explore new role in of SUBSrP in plants. Knock out of *OsSUBSrP1* produces apical abortion in KO-1, KO-2, and KO-3 verifying its indispensability for normal development in the rice.

In summary, our results revealed that SUBSrP1 is important to control the excessive outburst of ROS and programmed cell death (Figure 10). However, the mutation causes dysregulation of function in SUBSrP1 and excessive ROS induces abnormal cell death to apical spikelet of *apa1331.* In WT, the homeostasis in ROS and antioxidants produced a normal degeneration of tapetum that is essential for normal microspore development. However, excessive ROS and abnormal cell death cause the abnormal development of PMCs and lack of cuticle formation in *apa1331.* Mutation in *OsSUBSrP1* results in the impairment in transferring fatty acid molecules to the pollen development that causes sterility. Abnormal cutin formation and a significant decrease in wax and cutin biosynthesis cause the excessive water loss that produced aborted apical spikelet in *apa1331.* However, in WT, normal development of anther cuticle and wax saves water for cell viability and provides enough fatty acid molecules to develop pollen grains. Hence, our hypothesized model supports that SUBSrP1 is essential for maintaining ROS homeostasis and regular wax and cutin development in rice. Mutation in *OsSUBSrP1* produced the outburst of ROS and lack of anther cuticle formation due to downregulation of wax and cutin biosynthesis pathway genes that ultimately produce aborted apical spikelet in *apa1331*.

**Figure 10.**
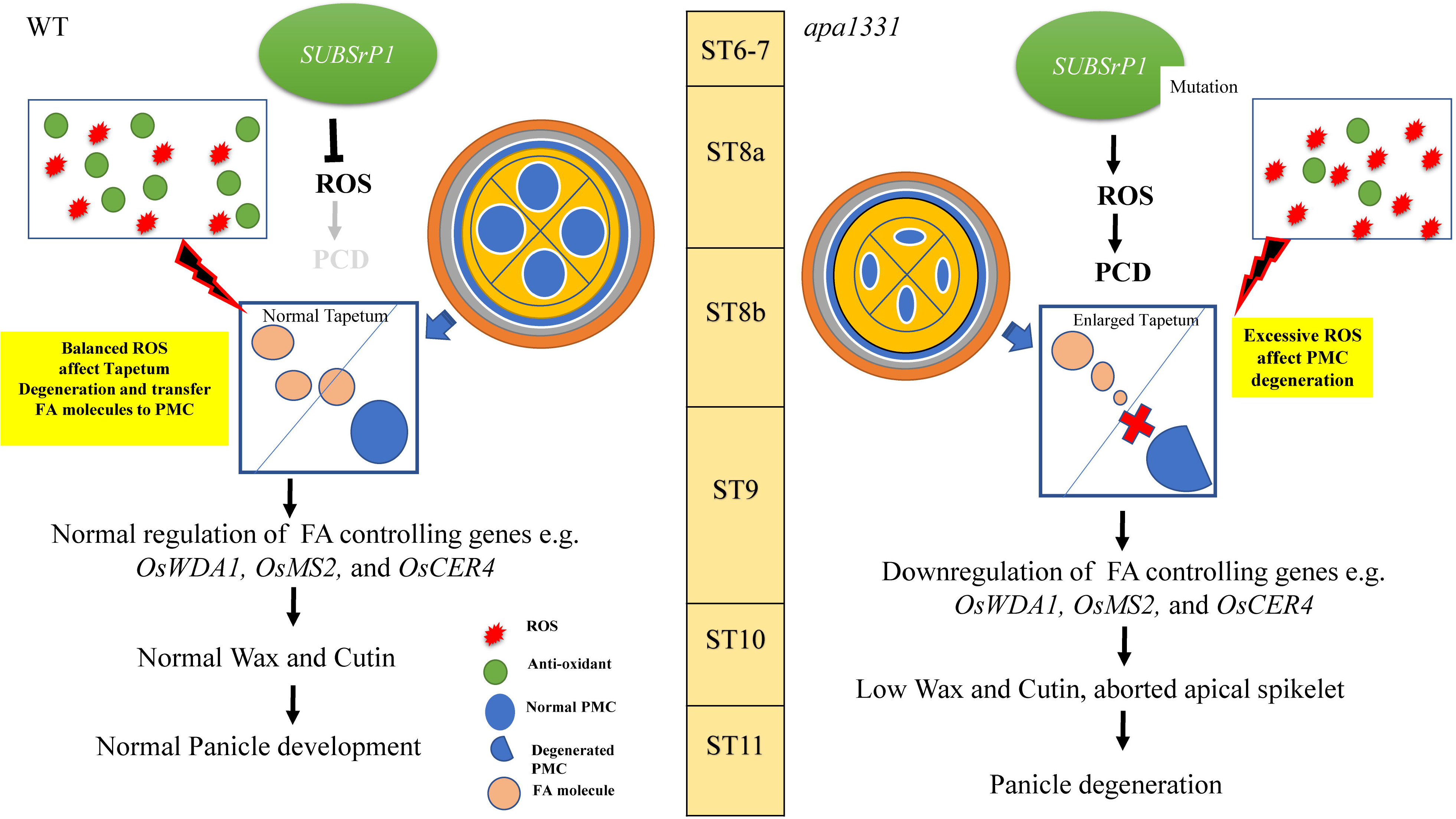
Illustration showing *SUBTILISIN-LIKE SERINE PROTEASE 1 (OsSUBSrP1)* homeostasis of Reactive oxygen species (ROS)-mediatad programmed cell death (PCD) for anther cuticle formation and seed setting rate. *OsSUBSrP1* is essential for maintaining ROS-mediated PCD and play a role the in the development of apical spikelet by regulating the expression of wax and cutin biosynthesis genes. A mutation causes the dysregulation of SUBSrP1, and excessive ROS caused an imbalance of homeostasis that caused a significant decrease in anther cuticle and defects in microspore development in *apical panicle abortion1331* (*apa1331).* Significant downregulation of fatty acid controlling genes caused a substantial reduction in wax and cutin content in *apa1331* compared to WT. The significant decrease of wax and cutin content in anthers of *apa1331* causes excessive water loss and cell death that causes abortion in apical spikelets.

## Acknowledgments

This work was supported by the National Key Research and Development of China (2016YFD0100406), Science and Technology Support Program of Sichuan (2017NZZJ005) and the National Natural Science Foundation of China (Grant No: 31771763). Authors are thankful to Dr. Hongyu Zhang for providing mutant for follow up functional study.

## Author contributions

AA and HZ, XW designed and conceived the experiments. AA and WTK performed the major experiments. PX and SAZ helped in the presentation of results and writing. AA wrote the manuscript. HZ, XQ, YL and LT helped in rice breeding and provided technical help during the study. XW and WW supervised the whole study with valuable suggestions. All authors read and approved the final version of manuscript.

## Data availability statement

The data supporting these findings are available upon request from the corresponding author Prof. Xianjun Wu.

## Table Legends

**Table S1.** List of primers used in the determination of relative expression

**Table S2.** List of primers used to validate the up-regulated genes

**Table S3.** List of primers used to validate the down-regulated genes

## Figure legends

**Figure 1S.** Validation of transcriptome revealed up and downregulated (differentially expressed genes) DEGs using qRT-PCR analysis

**(A)** Relative expression of up-regulated DEGs in *apical panicle abortion1331 (apa1331)* compared to Wild type (WT). **(B)** Relative of expression of downregulated DEGs in *apa1331* compared to WT. In **A-B** x-axis shows relative expression in WT and *apa1331* in blue and yellow color, respectively. In comparison, the y-axis shows the relative expression calculated of specific genes by qPCR.

